# HD-ZIP IV genes are essential for embryo initial cell polarization in the radial axis initiation in *Arabidopsis*

**DOI:** 10.1101/2024.04.27.588165

**Authors:** Sayuri Tanaka, Yuuki Matsushita, Yuga Hanaki, Takumi Higaki, Naoya Kamamoto, Katsuyoshi Matsushita, Tetsuya Higashiyama, Koichi Fujimoto, Minako Ueda

## Abstract

Plants develop along apical–basal and radial axes. In *Arabidopsis thaliana*, the radial axis becomes evident when the cells of the eight-cell proembryo divide periclinally, forming inner and outer cell layers. Although changes in cell polarity or morphology likely precede this oriented cell division, the initial events and the factors regulating radial axis formation remain elusive. Here, we report that three transcription factors belonging to class IV homeodomain-leucine zipper (HD-ZIP IV) family redundantly regulate radial pattern formation: HOMEODOMAIN GLABROUS11 (HDG11), HDG12, and PROTODERMAL FACTOR2 (PDF2). The *hdg11 hdg12 pdf2* triple mutant failed to undergo periclinal division at the eight-cell stage and cell differentiation along the radial axis. Live-cell imaging revealed that this failure in radial axis formation can be traced back to the behavior of the embryo initial cell (apical cell), which is generated by zygote division. In the wild type, the apical cell grows longitudinally and then radially and its nucleus remains at the bottom of the cell, where the vertical cell plate emerges. By contrast, the mutant apical cell elongates longitudinally and its nucleus releases from its basal position, resulting in a transverse division. Computer simulations based on the live-cell imaging data confirmed the importance of the geometric rule (the minimal plane principle and nucleus-passing principle) in determining the cell division plane. We propose that HDG11, HDG12, and PDF2 promote apical cell polarization, i.e., radial cell growth and basal nuclear retention, as the initial event of radial axis formation during embryogenesis.

## Introduction

Initiation of the main body axis is one of the first steps in developing an organism from a unicellular zygote. As the basic plant body is based on cylindrical structures, its patterning can be described using two axes; the apical–basal (shoot–root) and radial (inner–outer) axes. In *Arabidopsis* (*Arabidopsis thaliana*), the apical–basal axis is defined by the transverse cell division of the zygote, which generates an embryo initial apical cell and an extra-embryonic basal cell (1). The apical cell undergoes two rounds of vertical divisions (two- to four-cell stage embryo) and one round of transverse division (eight-cell stage embryo). The resulting spherical proembryo goes through one periclinal division that generates the inner and outer cell layers, defining the radial axis of the 16-cell stage embryo (2). The inner cell mass then specifies the vascular and ground tissues, while the outer layer differentiates into the protoderm (epidermis precursor) during subsequent embryogenesis (3). Live-cell imaging has revealed that dynamic cellular changes are the driving force of zygote polarization along the apical–basal axis (4-7). For example, the zygote elongates longitudinally in a manner of tip growth, and the nucleus migrates to the apical cell tip to establish the asymmetric cell division site (8, 9). By contrast, how and when the inner–outer polarization is initiated in the radial axis are still unknown. Furthermore, the molecular mechanisms underlying the radial axis formation are also obscure, whereas several key factors regulating apical–basal axis have been identified.

HOMEODOMAIN GLABROUS11 (HDG11) and HDG12 were identified as maternally derived transcription factors that regulate apical–basal axis formation (10). HDG11 and HDG12 (HDG11/12) and the paternally activated transcription factor WRKY2 all bind to the promoter of *WUSCHEL HOMEOBOX8* (*WOX8*) and initiate its *de novo* transcription after fertilization to trigger zygote polarization (10). After the zygotic cell division, *HDG11/12* expression is restricted to the basal cell derivatives (suspensor), thus maintaining *WOX8* expression in the suspensor (10). *HDG11/12* are also expressed in the embryo protoderm and post-embryonic epidermis (10-12). *WOX8* is not expressed in these tissues, suggesting as yet unidentified roles for HDG11/12 besides apical–basal axis formation.

*HDG11/12* belong to the class IV homeodomain-leucine zipper (HD-ZIP IV) gene family, several members of which are expressed in embryos, including the well-known epidermis-specific gene *ARABIDOPSIS THALIANA MERISTEM LAYER1* (*AtML1*) (12, 13). Similar to *HDG11/12, AtML1* is also expressed in the one-cell stage embryo and protoderm (13), but its role at such early stages is still unclear because the *atml1* mutant exhibits defects in epidermis morphogenesis only after the globular stage, even in a double mutant with its closest homolog, *PROTODERMAL FACTOR2* (*PDF2*) (14).

In this study, we demonstrate that these embryonic HD-ZIP IV genes play crucial roles in radial pattern formation, including proper cell differentiation. Furthermore, our live-cell imaging showed that these genes are necessary for the apical cell to start radial cell expansion. Based on computer simulations, we confirmed that the geometry of the apical cell and the position of its nucleus are important for determining the proper vertical division plane. Our findings show that HD-ZIP IV genes are crucial to set the radial axis initiation during very early embryogenesis.

## Results

### *HDG11/12, AtML1*, and *PDF2* show similar expression pattern during embryogenesis

We analyzed the expression patterns of *HDG11, HDG12, AtML1*, and *PDF2* using fluorescent reporter genes driven by each promoter, from the unfertilized egg cell to heart stage embryos (Fig. 1*A–D*). As we previously reported, we detected *HDG11*/*12* expression as early as in the egg cell (Fig. 1*A, B*) (10), and their expression remained through two- to four-cell stage embryos and finally became restricted to the suspensor and protoderm after the globular stage. The expression patterns of *AtML1* and *PDF2* were largely similar to those of *HDG11/12*, although unlike *HDG11*/*12*, the genes were not detectable in egg cells (Fig. 1*C, D*). The expression of these genes in egg cells and zygotes was consistent with published transcriptome data of isolated cells (15), supporting the idea that *HDG11/12, AtML1*, and *PDF2* all work during embryogenesis already in the zygote.

**Figure 1.**
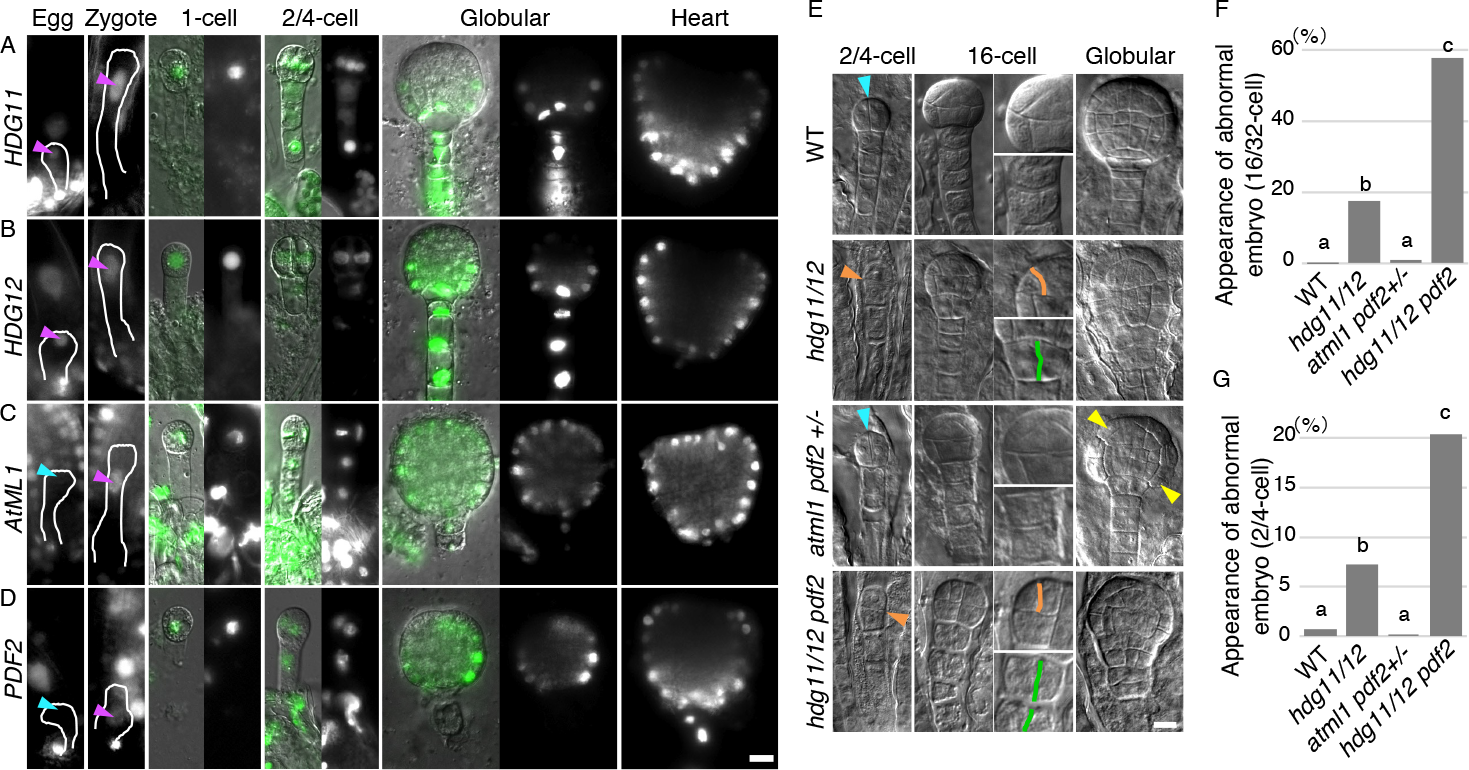
Expression patterns and roles of HDG11/12, *AtML1*, and *PDF2* on embryo patterning. (A–D) Epifluorescence microscopy images of the reporters *HDG11p::NLS-Venus* (A), *HDG12p::NLS-Venus* (B), *AtML1p::H2B-Clover* (C), and *PDF2p::H2B-Clover* (D) at the indicated stages of embryo development. The egg cells and zygotes are outlined. Magenta and cyan arrowheads indicate nuclei with and without fluorescent signals, respectively. The left panels of the one-cell, two- to four-cell, and globular stage embryo images were merged with differential interference contrast (DIC) images to show embryo structures. (E) DIC images of cleared embryos. Genotypes and embryo stages are indicated. Cyan and orange arrowheads at the two- to four-cell stage indicate proper (vertical) and abnormal (transverse) cell division planes of the apical cells, respectively. Right panels of 16-cell stage embryo images are enlarged images of the proembryo (top) and top part of suspensor (lower). Orange and green lines trace the aberrant cell division planes in the proembryo and suspensor, respectively. Yellow arrowheads in globular embryos indicate swollen epidermal cells. (F and G) Percentage of abnormal embryos, whose cell division patterns are different from those of the wild type (WT) at the 16- to 32-cell (F) and two- to four-cell (G) stages. Different lowercase letters indicate significant differences as determined by a Tukey– Kramer test; *P* < 0.05; n ≥ 112 (B); n ≥ 138 (C). Scale bars: 10 μm.

### HDG11/12 and PDF2 redundantly regulate cell division patterns after the zygotic division

We assessed the roles of HDG11/12, AtML1, and PDF2 in embryo formation by analyzing the phenotypes of their mutants (Fig. 1*E–G*). We previously reported that the double mutant *hdg11-1* and *hdg12-2* (*hdg11/12*), harboring loss-of-function T-DNA insertion mutants in *HDG11/12*, slightly affects the asymmetric zygote division and embryo shape at the globular stage (10). Here, we found that the uppermost suspensor cells divide vertically in *hdg11/12* at the 16-cell stage, as reported for the mutation of *WRKY2*, which acts together with *HDG11/12* on apical–basal axis formation (green line in Fig. 1*E*) (10, 16). In addition, *hdg11/12* embryos failed to properly establish the periclinal cell division that normally generates inner and outer cell layers (orange line in Fig. 1*E*). We observed aberrant cell division in the proembryo already at the two- to four-cell stage, as indicated by the transverse division of the apical cell (orange arrowhead in Fig. 1*E*). These abnormal cell divisions at both stages were significantly enhanced when the *pdf2-4* mutant was introduced into the *hdg11/12* mutant (*hdg11/12 pdf2*), although *pdf2-4* mutant did not show any cell division defects even when combined to *atml1-3* mutant, lacking function for its closely related gene, *AtML1* (*atml1 pdf2*+/-) (Fig. 1*E–G*).

The *atml1-3* and *pdf2-4* are both known as null alleles, and indeed, about 25% of the seeds produced in *atml1 pdf2*+/- plants contained arrested globular embryos harboring swollen epidermal cells as reported previously (yellow arrowheads in Fig. 1*E*) (14). The *hdg11-1 hdg12-2 atml1-3* triple mutant (*hdg11/12 atml1*) also showed abnormal cell division planes (Fig. S1*A*), but this genotype could not be used for reliable statistical analysis because of its low fertility (Fig. S1*B*). As an alternative, we therefore tested all possible double mutant combinations between *hdg11-1, hdg12-2, pdf2-4*, and *atml1-3*, which revealed that *pdf2-4* exhibits a slight but significant cell division defect when combined with either *hdg11-1* or *hdg12-2* (Fig. S1*C*). Another *PDF2* null allele, *pdf2-3*, showed the same effect as *pdf2-4* (Fig. S1*A*, compare to Fig. 1*E*). Furthermore, expression of *HDG11* or *PDF2* from the endogenous *HDG11* promoter partially rescued the cell division defects of *hdg11/12* embryos to similar levels (Fig. S2). These results demonstrate the redundant contribution of HDG11/12 and PDF2 to the regulation of cell division orientation already after zygote cell division.

### *PDF2* does not contribute to the apical–basal axis formation

Because HDG11/12 were identified as apical–basal axis regulators (10), we asked whether *PDF2* also participated in apical–basal axis formation, which starts with zygote polarization. The *hdg11/12* double mutant failed to elongate the zygote and to asymmetrically divide (Fig. 2*A*), as confirmed by the smaller total lengths of one-cell-stage embryos (Fig. 2*B*) and the higher ratio of apical divided by basal cell length (Fig. 2*C*). The *hdg11/12 pdf2* triple mutant also showed these defects to the same extent as *hdg11/12* (Fig. 2*A–C*).

**Figure 2.**
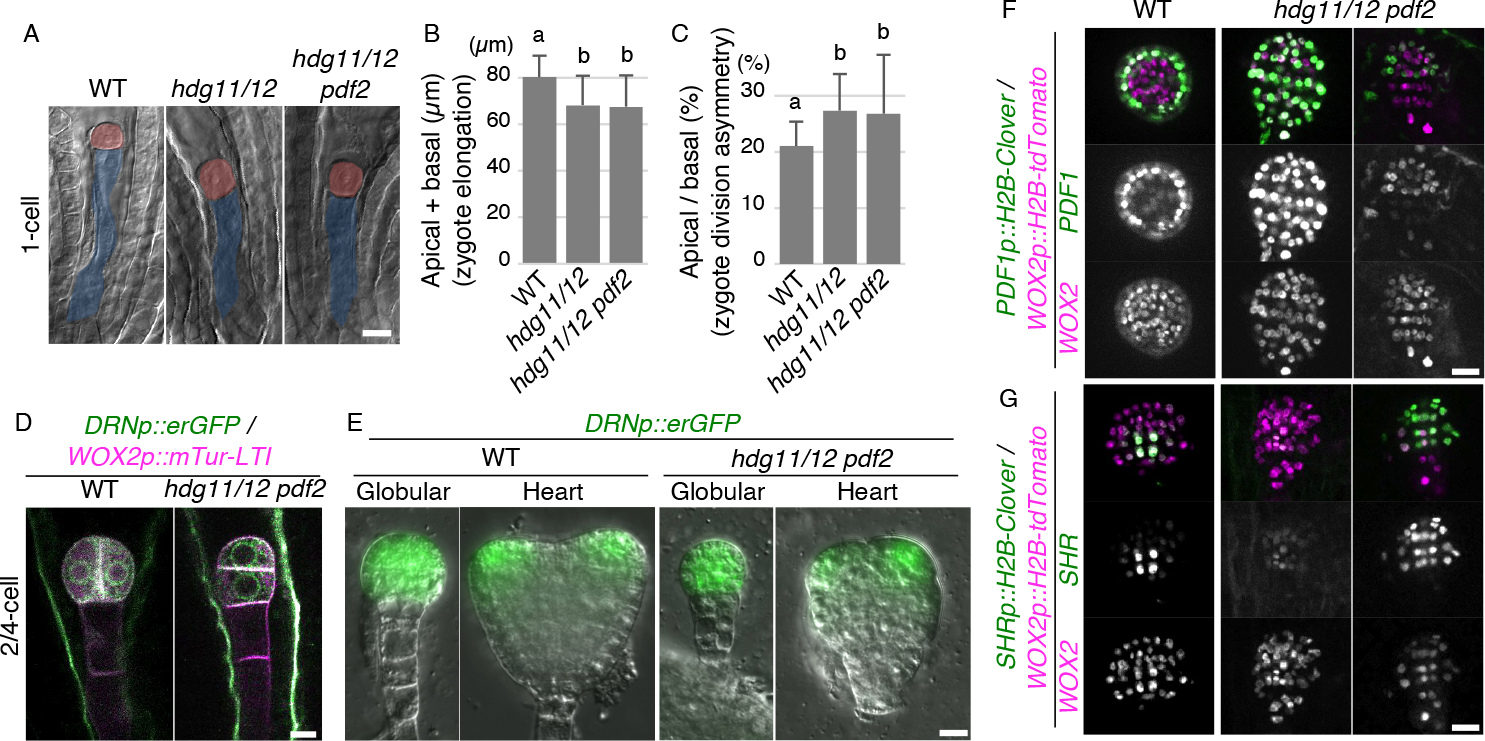
Roles of HDG11/12 and PDF2 in apical–basal and radial patterning. (A) DIC images of cleared one-cell stage embryos for the indicated genotypes. The apical (red) and basal (blue) cells are indicated. (B) Total length of one-cell stage embryos (sum of apical and basal cell lengths), denoted as zygote elongation. (C) Ratio of apical cell length divided by basal cell length in one-cell stage embryos, denoted as the asymmetric cell division of the zygote. Values are means ± standard deviation (SD, n = 80 for each genotype); different lowercase letters indicate significant differences as determined by a Tukey–Kramer test; *P* < 0.01. (D) Two-photon excitation microscopy images of the proembryo-specific marker *DRNp::erGFP* merged with the images of the embryo plasma membrane (PM) marker *WOX2p::mTur-LTI*. Maximum-intensity projection (MIP) images are shown. (E) Epifluorescence microscopy images of *DRNp::erGFP*. DIC images were merged to show embryo structures. (F and G) Confocal images of the epidermis-specific marker *PDF1p::H2B-Clover* (F) and the stele-specific marker *SHRp::H2B-Clover* (G) merged with the embryo nuclear marker *WOX2p::H2B-tdTomato* at the globular stage. Left and right panels of *hdg11/12 pdf2* show two different types of abnormal embryos. Lower panels show the enhanced images of individual fluorescence. Scale bars, 10 μm.

We also tested cell differentiation along the apical–basal axis using the proembryo-specific marker gene *DORNROSCHEN* (*DRNp::erGFP*) (17) and the suspensor-specific marker gene *WOX8* (*WOX8delta-Venus*) (16, 18). The *hdg11/12 pdf2* retained proper *DRNp::erGFP* signals in proembryos at all tested stages, although it showed aberrant cell division patterns (Fig. 2*D, E*). Similarly, *hdg11/12* and *hdg11/12 pdf2* displayed similar defects for *WOX8delta-Venus* expression, which was lost in the uppermost suspensor cells (Fig. S3*A*), as reported in *wrky2* mutant (16). These results suggest that PDF2 has no significant effect on the apical–basal axis.

### HDG11/12 and PDF2 are required for radial pattern formation

We focused on radial patterning because *hdg11/12 pdf2* failed to set proper periclinal cell division that separates inner and outer cell layers (Fig. 1*E*), which is defined as the initial event of radial axis formation (2). We tested the specification of radial tissues at the globular stage by generating the epidermis-specific markers for *PROTODERMAL FACTOR1* (*PDF1p::H2B-Clover*) and *AtML1* (*AtML1p::H2B-Clover*), as well as the vascular-specific marker for *SHORT-ROOT* (*SHRp::H2B-Clover*) (13, 19, 20). In the wild type, the expression of *PDF1p::H2B-Clover* and *AtML1p::H2B-Clover* was restricted to the outermost cell layers (Fig. 2*F* and Fig. S3*B*). By contrast, in *hdg11/12 pdf2*, we observed various patterns of expression; some embryos showed signals in entire cells, while others had no fluorescence in the outermost cells (Fig. 2*F* and Fig. S3*B*). Similarly, the *SHRp::H2B-Clover* expression domain was smaller or enlarged in *hdg11/12 pdf2* embryos (Fig. 2*G*). Together with the failure of periclinal divisions at the 16-cell stage, the observed defective inner–outer cell specification indicates that HDG11/12 and PDF2 are key factors in radial axis formation.

### HDG11/12 and PDF2 are required for radial cell growth and basal nuclear retention in the apical cell

We wished to identify the initial event regulated by HDG11/12 and PDF2. Because *hdg11/12 pdf2* was indistinguishable from *hdg11/12* for zygotic division (Fig. 2*A–C*), but enhanced the cell division defects of *hdg11/12* at the two- to four-cell stage (Fig. 1*G*), we checked when the radial axis becomes evident from the zygote to the 16-cell stage. To monitor the morphological changes and cell division patterns, we performed live-cell imaging with the marker *WOX2p::H2B–GFP* to label the embryo nucleus and *WOX2p::tdTomato-LTI6b* to label the plasma membrane (PM) (21, 22). In the wild type, we noticed that after the zygote division (yellow arrowheads in Fig. 3*A*, and Movie S1), the apical cell expands to take on a spherical cell shape before dividing vertically (cyan arrowhead in Fig. 3*A*). The resulting proembryo continued its oriented cell division and finally underwent periclinal divisions to form the inner and outer cell layers. By contrast, in *hdg11/12 pdf2*, even after the wild-type-like asymmetric zygote division, the apical cell elongated longitudinally and then divided transversely (orange arrowhead in Fig. 3*B*, and Movie S2). The resulting proembryo formed anticlinal cell divisions, failing to establish the inner–outer separation (magenta arrowhead in Fig. 3*B*).

**Figure 3.**
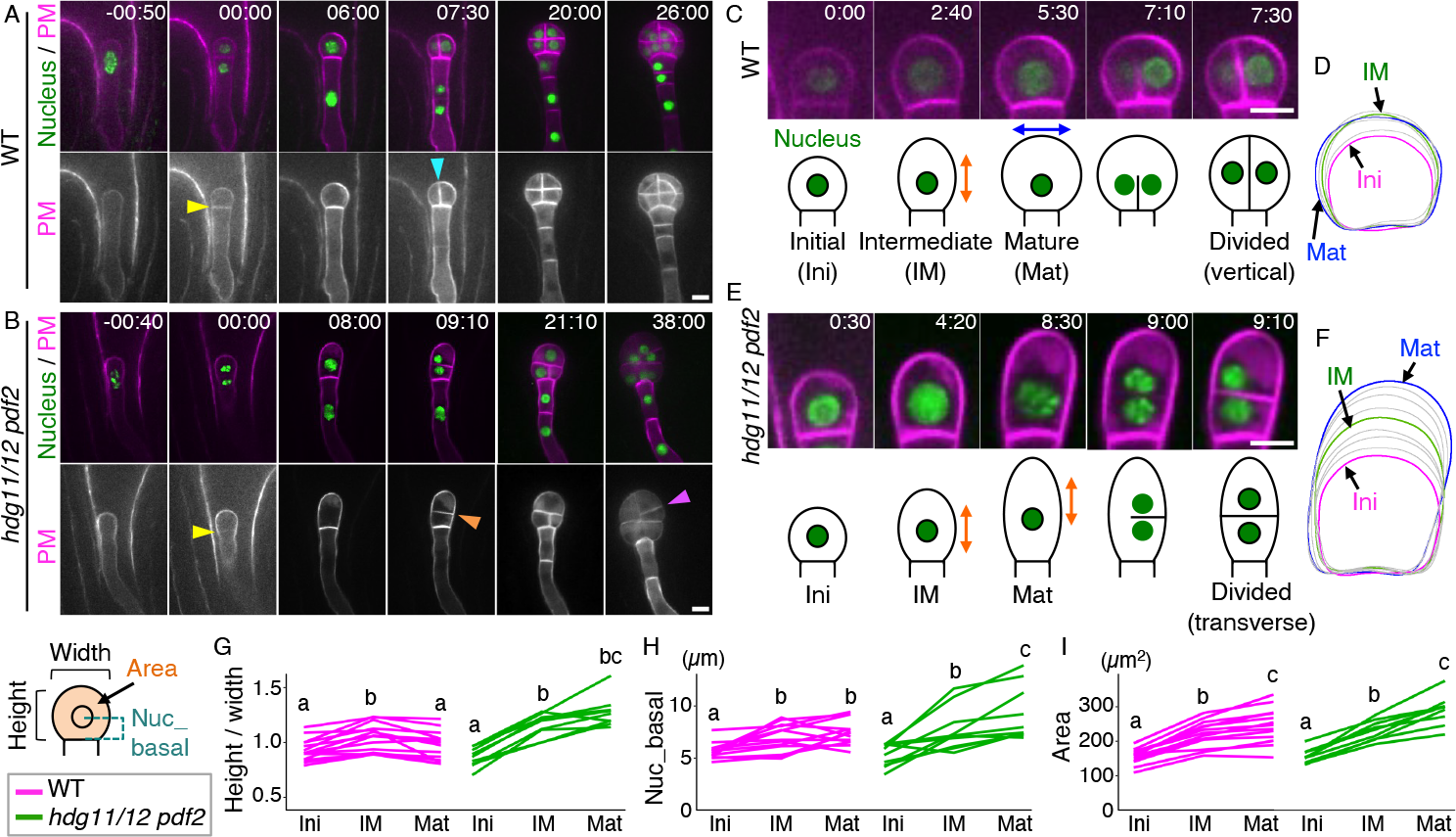
Live-cell imaging of radial pattern formation from the zygote. (A and B) Time-lapse observations of *in vitro*–cultivated WT (A) and *hdg11/12 pdf2* (B) ovules, expressing the embryo nuclear marker *WOX2p::H2B-GFP* and the PM marker *WOX2p::LTI-tdTomato*. Lower panels show enhanced PM images. Numbers indicate the time (h:min) from zygote division, as indicated by yellow arrowheads. Cyan and orange arrowheads indicate the vertical and transverse cell division planes, respectively. The magenta arrowhead indicates an abnormal anticlinal cell division plane. (C–F) Enlarged images of the apical cells (C and E) from the embryos shown in A and B, and the overlay of their sequential cell contour images (D and F). Lower panels show a summary of the respective stages, and cyan and orange arrows show the radial expansion and longitudinal elongation of the apical cell, respectively. The initial (Ini) cell is the cell with a clear boundary between apical and basal cell after zygote division; the mature cell (Mat) is just before nuclear division. Intermediate (IM) is the mid-time point between Ini and Mat; the divided cell is just after the completion of cell plate formation. The cell contours of these time points are colored in D and F. (G–I) Graphs of the ratio of apical cell height divided by apical cell width (height/width; G), distance between the nuclear center and the cell bottom (H), and apical cell area (I) at the indicated stages as shown in the left illustration. Different lowercase letters indicate significant differences, as determined by a Wilcoxon signed-rank test with Bonferroni correction for multiple samples; *P* < 0.05. WT, n = 14 (G and I) and n = 13 (H); *hdg11/12 pdf2*, n = 9 (G and I) and n = 10 (H). Scale bars, 10 μm.

Further focusing on the behavior of the apical cell revealed that the wild-type cell elongates slightly longitudinally and then expands radially (Fig. 3*C*), as shown by the overlay of sequential cell contour images (Fig. 3*D*). The cell plate finally emerged vertically from the cell bottom and reached the cell tip. We detected this polar cell plate emergence in all observed wild type embryos (100%, n = 18; Movie S3). By contrast, the apical cell of *hdg11/12 pdf2* continued to elongate longitudinally, and a transverse cell plate formed (Fig. 3*E, F*). As shown in Fig 1*G*, the transverse division appeared in about 20% of the total mutant embryos, and the cell plate emerged from the lateral side in most of these embryos (79%, n = 14; Movie S4). We confirmed the alteration of growth direction in the wild type and the continuous elongation in the mutant by calculating the ratio between cell height and cell width based on our live-cell imaging data (Fig. S4*A*) and the significant difference after zygote division (initial; Ini), just before apical cell division (mature; Mat), and their intermediate time point (IM) (Fig. 3*G*). Consistent with the cell plate emergence site, we also found that the nucleus is retained at the bottom region during apical cell growth in the wild type but not in the mutant (Fig. 3*H*). Notably, the apical cell area increased to a similar extent in the wild type and mutant (Fig. 3*I*, and Fig. S4*B*), indicating that *HDG11/12* and *PDF2* are specifically required for cell polarization, i.e., radial cell growth and basal nuclear retention.

### Apical cell geometry and nuclear position are sufficient to predict the cell division plane in wild type

Recent geometric simulations based on fixed samples showing divided sister cell shapes successfully inferred the cell division patterns of embryos from the one-cell to the 16-cell stage (23, 24). In these simulations, the cell division plane can be predicted based on two principles (geometric rule): 1) the new cell division plane should pass through the cell centroid, and 2) the plane area should be minimal. Moreover, the nuclear position might substitute for the cell centroid, as with leaf stomatal cell precursors where the nucleus polarly migrates to the cell corner to induce a marked asymmetric cell division (24, 25). Since this idea fits our findings of nuclear retention and cell plate emergence at the cell bottom in wild-type apical cells (Fig. 3*C*), we performed simulations using our time-lapse data to evaluate the contribution of the cell centroid and nucleus in determining the cell division plane. We first used wild-type data to extract two-dimensional cell contours and nucleus position from Mat cell images and inferred the cell division planes (Cellular Potts model; Fig. 4*A*). In this simulation, we comprehensively explored the cell division planes whose lengths should be small and whose inferred ratios to the sister cell area (*ρ*_*inf*_) should be close to the observed ratio (*ρ*_*obs*_). We then evaluated the simulated planes by comparing these values to the actual image of the divided cell stage for the same embryo, as well as to the centroid and nuclear position data of the Mat cell. In the simulated data, the length of the cell division plane was smaller in the vertical division planes than in the transverse planes (Fig. 4*B, C*). Furthermore, in the vertical planes, those with shorter distances to the centroid or nucleus showed *ρ*_*inf*_ values close to *ρ*_*obs*_ (magenta in Fig. 4*B* and *C*). The inferred area ratios based on the nucleus-passing principle (*ρ*_*inf*_ (nucleus)) matched *ρ*_*obs*_ better than those based on the centroid-passing principle (*ρ*_*inf*_ (centroid)) (Fig. 4*D, E*), as shown by the higher correlation coefficient (0.60 for *ρ*_*inf*_ (nucleus) and 0.15 for *ρ*_*inf*_ (centroid)). These results indicate that the cell geometry and nuclear position, rather than the cell centroid, are sufficient to predict the cell division plane in the wild type, supporting the importance of a basally retained nucleus in the apical cell.

**Figure 4.**
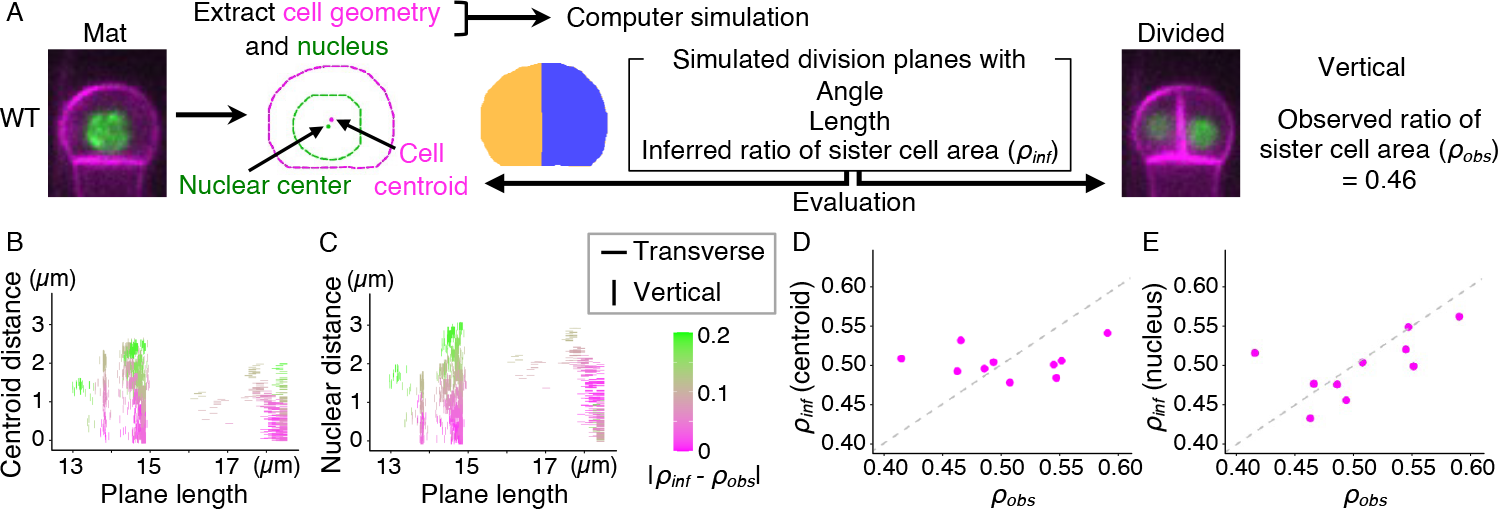
Computational evaluation of the geometric rule using time-lapse images before and after apical cell division. (A) Procedure of the image extraction, simulation, and evaluation using live-cell imaging data of WT embryos. The mature cell (Mat) is just before nuclear division, and the divided cell is just after the completion of cell plate formation, as indicated in Fig. 3. (B and C) Distribution of simulated cell division planes on the plane length and distance to cell centroid (B) or nuclear center (C). Simulated planes with vertical (90º ± 15) and transverse (0º ± 15) angles are plotted as vertical and transverse bars, respectively. The right color bar shows the absolute difference of the inferred ratio of sister cell area (*ρ*_*inf*_) from the observed ratio (*ρ*_*obs*_), which is set as the reference point (0; magenta). (D and E) Scatterplots exploring the relationship between *ρ*_*obs*_ and inferred area ratios based on centroid-passing principle *ρ*_*inf*_ (centroid) (D) and nucleus-passing principle *ρ*_*inf*_ (nucleus) (E). n = 10. The dotted lines represent *ρ*_*inf*_ = *ρ*_*obs*_.

### The *hdg11/12 pdf2* apical cells divide transversely by following the geometric rule

To determine whether *hdg11/12 pdf2* also follows the geometric rule, we performed the simulation using data for the mutant Mat apical cells (Fig. 5*A*). These simulations showed that the transverse planes are shorter in the mutant than the vertical planes (Fig. 5*B, C*), in contrast to the wild type (Fig. 4*B, C*). For the transverse planes, those with shorter distances to the centroid or nucleus showed inferred ratios of sister cell area (*ρ*_*inf*_) that are close to the observed ratio (*ρ*_*obs*_) (magenta in Fig. 4*B* and *C*). The predominance of nucleus- and centroid-passing principles on sister cell area determination was not clearly different, as shown by their similar correlation coefficient (0.68 of *ρ*_*inf*_ (nucleus) and 0.63 of *ρ*_*inf*_ (centroid)) (Fig. 5*D, E*), possibly consistent with the failure to retain the nucleus in a basal position. These results show that the apical cells of the *hdg11/12 pdf2* triple mutant follow the geometric rule, resulting in a transverse division due to abnormal cell shape and nuclear position.

**Figure 5.**
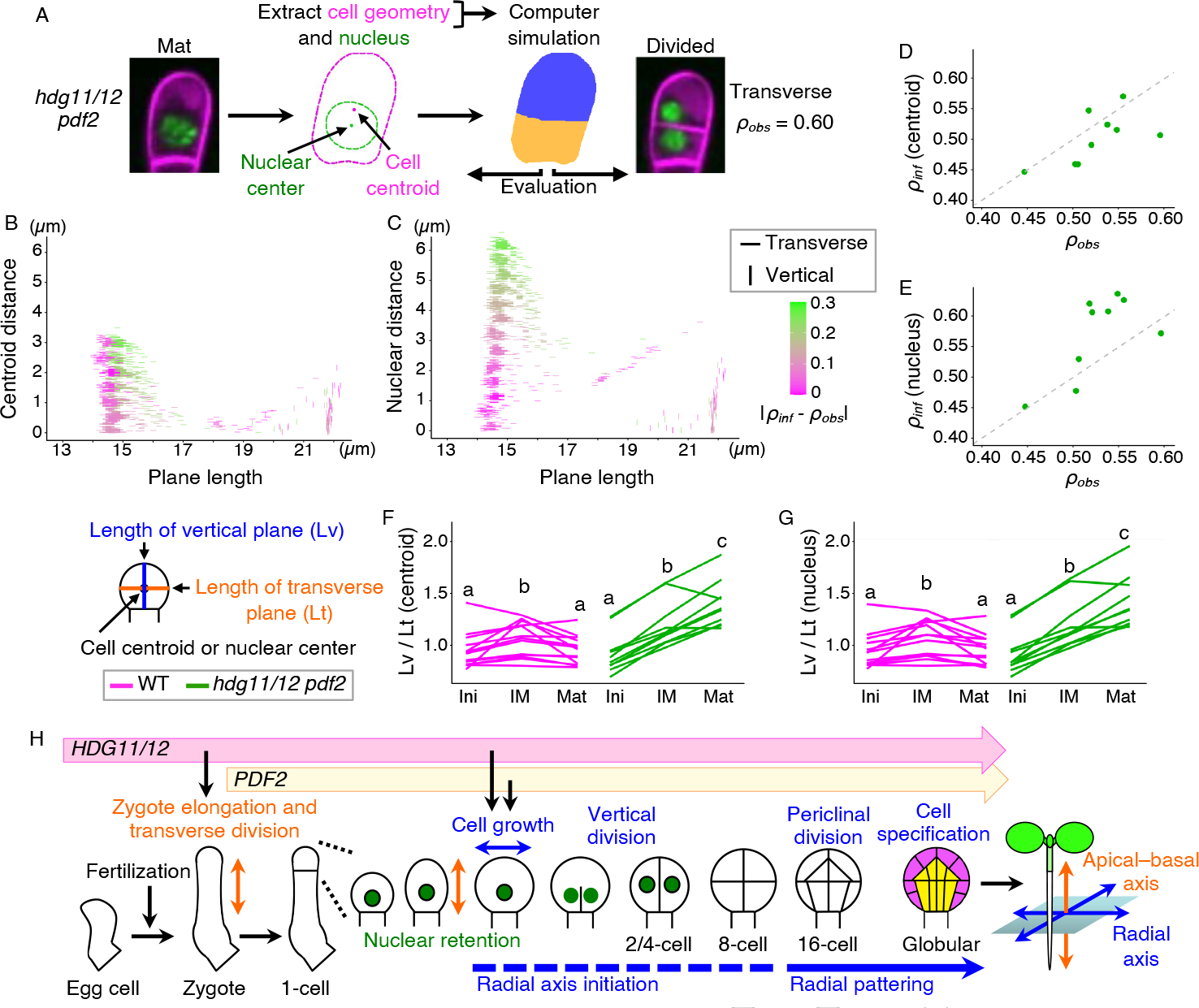
Computational evaluation of the geometric rule in *hdg11/12 pdf2*. (A) Procedure of the simulation analysis of *hdg11/12 pdf2* embryo. (B and C) Distribution of simulated cell division planes on the plane length and distance to cell centroid (B) and nuclear center (C). Simulated planes with vertical (90º ± 15) and transverse (0º ± 15) angles are plotted as vertical and transverse bars, respectively. The right color bar shows the absolute difference of *ρ*_*inf*_ from *ρ*_*obs*_, which is set as the reference point (0; magenta). (D and E) Scatterplots exploring the relationship between *ρ*_*obs*_ and inferred area ratios *ρ*_*inf*_ (centroid) (D) and *ρ*_*inf*_ (nucleus) (E). n = 9. The dotted lines represent *ρ*_*inf*_ = *ρ*_*obs*_. (F and G) Ratio of the vertical plane length (Lv) divided by the length of the transverse plane (Lt), both of which pass through the centroid (F) or nucleus (G) at the indicated stages as shown in the left illustration. Different lowercase letters indicate significant differences among the three stages, as determined by a Wilcoxon signed-rank test with Bonferroni correction for multiple samples; *P* < 0.05. WT, n = 13; *hdg11/12 pdf2*, n = 10. (H) Diagram summarizing the expression period of HD-ZIP IV genes and the dynamics of radial axis formation.

### Preparing proper cell polarity during apical cell maturation is important to select the proper cell division plane

Since cell growth direction and nuclear position after the IM stage differed between the wild type and *hdg11/12 pdf2* (Fig. 3), we investigated when the geometric conditions suitable for proper cell division are met. We thus performed geometric simulations at three time points, Ini, IM, and Mat in the wild type (Fig. S5*A–C*). In both cases of the principle passing through the nucleus and centroid, the simulated planes that followed the geometric rule (i.e., the samples appearing in the lower left area of each plot in Fig. S5*B, C*) were the transverse planes showing a close area ratio *ρ*_*inf*_ to *ρ*_*obs*_ at the Ini and Mat stages. At the IM stage, however, the planes whose *ρ*_*inf*_ did not match *ρ*_*obs*_ followed the geometric rule, possibly due to immature cell shape and fluctuations in nuclear position. This result suggested that the geometric condition in the IM stage may not be sufficient to induce the formation of a proper cell division plane. Thus, we compared the length of the vertical plane (Lv) and transverse plane (Lt), which pass through the nucleus or centroid at each stage (Fig. 5*F, G*). In the wild type, the ratio of Lv to Lt was significantly lower in the Mat stage after a transient increase in the IM stage in both cases of nucleus and centroid, suggesting that the tendency of selecting the proper transverse division plane increases in the Mat stage based on the geometric rule.

In *hdg11/12 pdf2*, at the Ini stage, the simulated planes that followed the geometric rule were the vertical planes showing *ρ*_*inf*_ values close to *ρ*_*obs*_ as in the wild type (Fig. S6, compare to Fig. S5). However, the planes that followed the geometric rule changed to the transverse planes at the IM stage, with most of the simulated results finally showing the transverse planes being those following the geometric rule at the Mat stage. In agreement with this observation, the ratio of Lv to Lt significantly increased with time in *hdg11/12 pdf2* (Fig. 5*F, G*), showing a gradual increase in the predominance of transverse division based on the geometric rule.

We conclude that cell polarization during apical cell maturation, i.e., initiation of radial growth and basal nuclear retention, is important for selecting the proper vertical cell division plane based on the geometric rule and that *hdg11/12 pdf2* fails to set this polarity, resulting in transverse cell divisions.

## Discussion

Our genetic analysis revealed that HDG11/12 and PDF2 are key regulators of radial patterning during embryogenesis. Furthermore, by combining live-cell imaging, image quantification, and computer simulations, we demonstrated that these HD-ZIP IV genes are crucial for radial cell growth and basal nuclear retention in the apical cell, both necessary steps in determining the manner of cell division based on the geometric rule. These findings provide the intriguing model that the radial axis is already initiated just after the asymmetric zygote division that defines the apical–basal axis (Fig. 5*H*).

We found that HDG11/12 and PDF2 redundantly regulate radial patterning and that *PDF2* expression can partially rescue the *hdg11/12* defects. Nevertheless, PDF2 had no detectable effect on apical–basal axis formation, in which HDG11/12 play important roles as maternally derived factors to the zygote (10). One possible explanation for these different contributions is their different expression levels in the egg cell, i.e., the expression of *HDG11/12* and the lack of *PDF2* expression (Fig. 1). Because the transgenic heterologous expression of *HDG11* in the sperm cell was sufficient to fully rescue the *hdg11/12* zygote defects, HDG11/12 stored before fertilization may be crucial to initiate the formation of the apical–basal axis after fertilization (10). Therefore, *PDF2* expression from the zygote, together with already present HDG11/12, would regulate apical cell activity (Fig. 5*H*).

Our quantitative live-cell imaging revealed that HDG11/12 and PDF2 are required for radial cell growth and basal nuclear retention in the apical cell (Fig. 3). Computational simulations showed the direct contribution of these two parameters in determining the proper cell division plane, confirming the importance of the geometric rule, i.e., the minimal plane principle and the nucleus-passing principle (Figs. 4 and 5). The predominance of the nucleus-passing principle over the centroid-passing principle was previously demonstrated by simulations using stomatal cell precursors, which take on an asymmetric cell shape and undergo long-distance nuclear migration (24, 25). The apical cell division is not asymmetric but requires strict directional control to initiate pattern formation. Our findings reveal the subtle but important polarization prior to cell division timing that ensures proper cell division.

The longitudinal elongation of the initial apical cell resembles that of the zygote. Considering that the apical cell in the *hdg11/12 pdf2* triple mutant continues to elongate longitudinally, HD-ZIP IV genes may function to switch from the longitudinal polarity in the zygote to radial polarity in the apical cell (Fig. 5*H*). In addition, the nucleus was not retained at the cell bottom in *hdg11/12 pdf2*, and simulations for this mutant based on the nucleus- and centroid-passing principles were both correlated with the actual division pattern. Therefore, HD-ZIP IV genes likely polarly hold the nucleus away from the default position (the cell centroid) possibly via actin filament (F-actin) or microtubules (MT), which are the basis of polar nuclear positioning in zygotes or stomatal cell precursors, respectively (9, 25). Since plant cell growth is generally directed by cortical MTs, and previous simulations of early embryogenesis demonstrated the correlation between the cortical MT pattern and division plane orientation (26), the roles of cortical MTs in radial cell growth need to be assessed.

Since HDG11/12 act as transcriptional activators in the zygote (10), it is important to identify the genes acting downstream of HDG11/12 and PDF2 in the apical cell and the proembryo. A possible player may be the regulatory genes related to plant hormone auxin because HDG11 can indirectly activate auxin signaling during lateral root formation (27), and altered divisions of the apical cell and proembryo were reported in several mutants of auxin regulators, such as the auxin biosynthetic enzyme TRYPTOPHAN AMINOTRANSFERASE OF ARABIDOPSIS 1 (TAA1) and the auxin-responsive transcription factors MONOPTEROS (MP) and BODENLOS (BDL) (28-30). The auxin pathway affects the pattern of MTs and F-actin to set the orientation of periclinal division from the eight-cell to the 16-cell stage embryo (31). Notably, the pathways involving auxin and HD-ZIP IV genes do not fully overlap, as *hdg11/12 pdf2* showed proper expression of *DRN*, a direct target of MP (Fig. 2) (17); conversely, the *mp* mutant still retains proper *AtML1* expression in the proembryo epidermis, whereas *AtML1* expression was severely disrupted in *hdg11/12 pdf2* (Fig. S3) (13). *AtML1* expression is robustly maintained by various mechanisms, and not impaired in general embryo patterning mutants (13, 31-33), supporting the importance of HDG11/12 and PDF2 in radial axis initiation. Further combination of mutant analysis, quantitative live-cell imaging, and computational simulations will help reveal the molecular mechanism underlying the geometric rule and radial axis formation.

## Materials and Methods

### Mutant strains and growth conditions

All *Arabidopsis* lines were in the Columbia (Col-0) background. *hdg11-1* (SAIL_865 G09), *hdg12-2* (SALK_127261), *atml1-3* (SALK_033408), *pdf2-3* (GK-201A01), and *pdf2-4* (SAIL_70_G06) have been described previously (10, 12, 14). Plants were grown at 18–22°C under continuous light or long-day conditions (16-h-light/8-h-dark photoperiod).

### Reporter lines and plasmid construction

The following transgenic lines have been described previously: *HDG11p::NLS-Venus, HDG12p::NLS-Venus* (10), *WOX2p::LTI-tdTomato, WOX2p::H2B-GFP* (21), *WOX8delta-Venus* (16), and *DRNp::erGFP* (17). The fluorescent reporters for *AtML1, PDF2, PDF1*, and *SHR* contained promoter fragments of the respective genes driving the expression of *HISTONE H2B* (*H2B*) or the GFP-derived reporter gene *Clover*.

### Histological analysis and time-lapse observation

Differential interference contrast (DIC) microscopy observations of embryos were performed as previously described (10). The live-cell imaging of embryos using *in vitro* ovule cultivation was performed as in a previous report with some modifications (21, 34, 35).

Cell geometry, e.g., cell shape and nucleus (centroid) position, were extracted from experimental data with ImageJ (https://imagej.nih.gov/ij/index.html) and cellpose (36). These geometric data were then smoothed with Python to introduce geometric simulations. Based on the segmented images of the cells and nuclei, the following parameters were measured using ImageJ software: cell area, cell width (defined as the width of the cell bounding box), cell height (defined as the height of the cell bounding box), and distance between the nuclear center and the cell bottom.

### Computer simulations of apical cell division planes

In our geometric simulations (Figs. 4 and 5), candidates of cell division planes were numerically explored through the Cellular Potts Model, which gives the minimal area surface depending on cell shape (24, 37). The plane length and nucleus (centroid) distance of planes were individually evaluated and quantified from the results of Cellular Potts simulations. The division plane characteristics (e.g., cell area ratio and plane contour length ratio) were inferred by selecting division planes that obey geometric rules.

## Acknowledgments

We thank Kohdai Nakajima, Yusuke Kimata, Hikari Matsumoto, Yoshikatsu Sato, Zichen Kang, and Satoru Tsugawa for technical support, Taku Takahashi for providing *hdg11/12* double mutants, Wolfgang Werr for *DRNp::erGFP* and Thomas Laux for *WOX8delta-Venus*. Microscopy was conducted at the Institute of Transformative Bio-Molecules (WPI-ITbM) of Nagoya University. This work was supported by the Japan Society for the Promotion of Science [a Grant-in-Aid for Scientific Research on Innovative Areas (JP19H05670, and JP19H05676 to M.U.; JP22H04719 to K.F.; JP16H06280 (Advanced Bioimaging Support)), a Grant-in-Aid for Scientific Research (B) (JP23H02494 to M.U.; JP20H03289 to T.Higaki), and International Leading Research KEPLR (JP22K21352) to T.Higashiyama and M.U.], the Japan Science and Technology Agency [CREST (JPMJCR2121 to M.U., T.Higaki, and K.F.)], the Suntory Rising Stars Encouragement Program in Life Sciences (SunRiSE; to M.U.), and the Toray Science Foundation (20-6102 to M.U.).

