## Supplementary material for "HD-ZIP IV genes are essential for embryo initial cell polarization in the radial axis initiation in *Arabidopsis*": Fig. S

### Supplementary Materials and Methods

### Plasmid construction

The promoter lengths and construct codes for each promoter reporter are: *AtML1* (3.4 kb, MU1861), *PDF2* (2.4 kb, MU2303), *PDF1* (1.6 kb, MU1850), and *SHR* (2.8 kb, MU2251). Each promoter fragment was combined with the full-length coding region of histone 2B (H2B; AT1G07790), GFP-derived Clover, and the NOS terminator in a pPZP221 binary vector (1), only except for MU2251, where pMDC99 binary vector was used (2).

In *WOX2p::H2B-tdTomato* (MU1933), the 3.5-kb *WUS HOMEOBOX2* (*WOX2*; AT5G59340) promoter was fused to H2B-tdTomato and NOS terminator in the above pMDC99 binary vector.

The PM marker (*WOX2p::mTur-LTI*, MU2329) contains the *WOX2* promoter, cyan-fluorescent mTurquoise2, the full-length coding region of LOW TEMPERATURE INDUCED PROTEIN 6B (LTI6b, AT3G05890), and the NOS terminator in a pMDC100 binary vector (2).

In *HDG11p::HDG11-YFP* (MU2383) and *HDG11p::PDF2-YFP* (MU2388), the 2.6-kb *HDG11* promoter was fused to the full-length coding region of *HDG11* or *PDF2* with yellow-fluorescent Venus and 35S-terminator in pMDC99 binary vector, respectively.

These constructs were transformed into *Arabidopsis* using the floral dip method (3).

### Histological analysis and microscopy

The DIC observation of cleared seeds was performed using Nomarski microscope (AxioImager A2; Zeiss), as previously reported (4). Epi-fluorescent images of fluorescent reporters were also acquired using AxioImager A2. For confocal observation of tissue-specific reporters, the dissected embryos were observed using an inverted confocal microscope system (CV1000; Yokogawa Electric) as previously reported (5). The *in vitro* ovule cultivation and time-lapse imaging of embryo patterning was performed using a spinning-disk confocal microscope system (CSU-W1; Yokogawa Electric), as shown in a previous report with some modifications (5-7). The two-photon excitation microscopy images were acquired using a laser-scanning inverted microscope (A1R MP; Nikon) equipped with a Ti:sapphire femtosecond pulse laser (Mai Tai DeepSee; Spectra-Physics, Mountain View, CA, USA), as previously reported (8, 9).

### Computational simulation of apical cell division planes

We used Cellular Potts model to explore cell division plane with minimal surface. We defined site $i$ for each pixel which constitutes image extracted and smoothed from experimental data. For each site $i$, we assigned $x[i]$, as a variable attributing to either of daughter cells ($x[i] = 1{,}2$), explored the configurations which minimize the following energy function $\mathcal{H}$ by Monte Carlo simulation.

Following the previous study (10), we defined energy function $\mathcal{H}(x)$ as

$$\mathcal{H} = \mathcal{H}_V + \mathcal{H}_A.$$

$\mathcal{H}_V(x)$ is a penalty term to keep the daughter-cell areas to be targeted areas. In Cellular Potts simulation, actual cell areas of two daughter cells ($V_1, V_2$) can be represented as

$$V_1 = \sum_{\substack{i \text{ in} \\ \text{all sites}}} \mathbb{1}_{x[i]=1}, V_2 = \sum_{\substack{i \text{ in} \\ \text{all sites}}} \mathbb{1}_{x[i]=2},$$

where a summation is over all the sites constituting mother cell. $\mathbb{1}_{\text{condition}}$ is the indicator function whose value is assigned $1$ when the condition is satistified, and $0$ otherwise.

Then, $\mathcal{H}_V(x)$ can be written down as

$$\mathcal{H}_V(x) = (V_1 - V_1^*)^2 + (V_2 - V_2^*)^2,$$

where $V_1^*, V_2^*$ are targeted cell area of two daughter cells. Introducing target area ratio $\rho^*$ and mother cell area $V$, we can rewrite $V_1^*, V_2^*$ as

$$V_1^* = \rho^* V, V_2^* = (1 - \rho^*)V.$$

$\mathcal{H}_A(x)$ is a function, which penalizes the increase of interface are between two daughter cells, as

$$\mathcal{H}_A(x) = \sum_{\substack{(i,j) \text{ in} \\ \text{neighbors}}} \mathbb{1}_{x[i] \neq x[j]},$$

where we summed up twelve cells from first nearest neighbors to third nearest-neighbors lattice pairs $(i, j)$. In conventional simulations of cell sorting, Metropolis-Hasting method using a state copy is well used in the Monte Carlo simulation (11). However, the method using the copy is redundant for our purpose that is a sampling of sufficient low energy states. To avoid this redundancy, we used the following simple Metropolis method at boundary. This method consists of stochastic updates of $x[i]$. For each update, we chose single site $i$ and checked the whether the site $i$ located in the boundary between daughter cells. If it is located the boundary, we prepared a candidate state whose is $x[i]$ changed to the other state from the current state. Then, we computed the excess energy of the candidate state from the current state, $\Delta E$. We accepted the update to the candidate state with the probability $\exp(-\beta \Delta E)$, and rejected it otherwise. We defined consecutive $N$ updates as one Monte Carlo Step (MCS), where $N$ is the pixel number of the focal cell measured at the initial, intermediate or mature phase *in vivo*. We computed configurations which have minimal energy with various $\rho^* = 0.35, 0.375, \ldots, 0.475, 0.5$. 600 random initial configurations are prepared for each $\rho^*$, and we obtained resulting configurations after $1 \times 10^4$ MCS. For efficient relaxation, we scheduled Monte Carlo simulation into two equally separated schemes with $\beta = 0.5 \to 2$.

**Evaluation of simulated planes**

For evaluating the planes in the context of cell division process, we first excluded simulated configurations which have domains more than three, since this output is obviously out of our interest. Over remaining planes, we quantify the division pattern, plane length, and distance to nucleus (or centroid) as follows. We defined division patterns in *silico* as the angle between the basal cell wall and line passing through simulated daughter cells' centroid. We rotated experimental data so that the basal cell wall possessed at the bottom (as shown in Figs. 4 and 5), and then we computed the daughter cells' centroid and angle of the line passing through them. If the angle is $90°$ ($0°$), daughter cells are divided vertically (transversely). We defined the case with $90° \pm 15°$ as vertical division pattern, and $0° \pm 15°$ as horizontal division pattern.

Plane length can be calculated as

$$(\text{plane length}) = \frac{1}{2} \sum_{\substack{(i,j) \text{ in} \\ \text{nearest neighbors}}} \mathbb{1}_{x[i] \neq x[j]}.$$

By calculating nucleus (or centroid) position in lattice space, we defined the distance to nucleus (or centroid) from plane as their minimal distance.

Here, we note that the case with $\rho^* \neq 0.5$, i.e., asymmetrical division case, we need to distinguish which of daughter cells is larger. For example, if we observe asymmetric transverse division pattern with $\rho^* = 0.4$, two configurations are possible; bottom daughter cell is larger or top daughter cell is larger. This difference is biologcally essential. Thus, we re-labeled $\rho^*$ value to $1 - \rho^*$ when top cell is larger in transverse division case, as well as left cell is larger in vertical division case.

**Inferring division plane characteristics from simulated planes**

All the explored planes are characterized by their orientation (i.e., vertical or transverse), plane length, distance to nucleus or cell centroid, divided cell area ratio $\rho^*$. Then, we selected planes by the following steps, as adopting geometric rules. (i) We selected 40% of planes which has less distance to nucleus (centroid) of all the simulated planes. (ii) Among the 40% we selected, we further selected 50% planes which has less plane length. We inferred the characteristic values, the cell area ratio $\rho_{inf}$ (nucleus) ($\rho_{inf}$ (centroid)) or plane length (Lv, Lt) as their ensemble average among the selected planes.

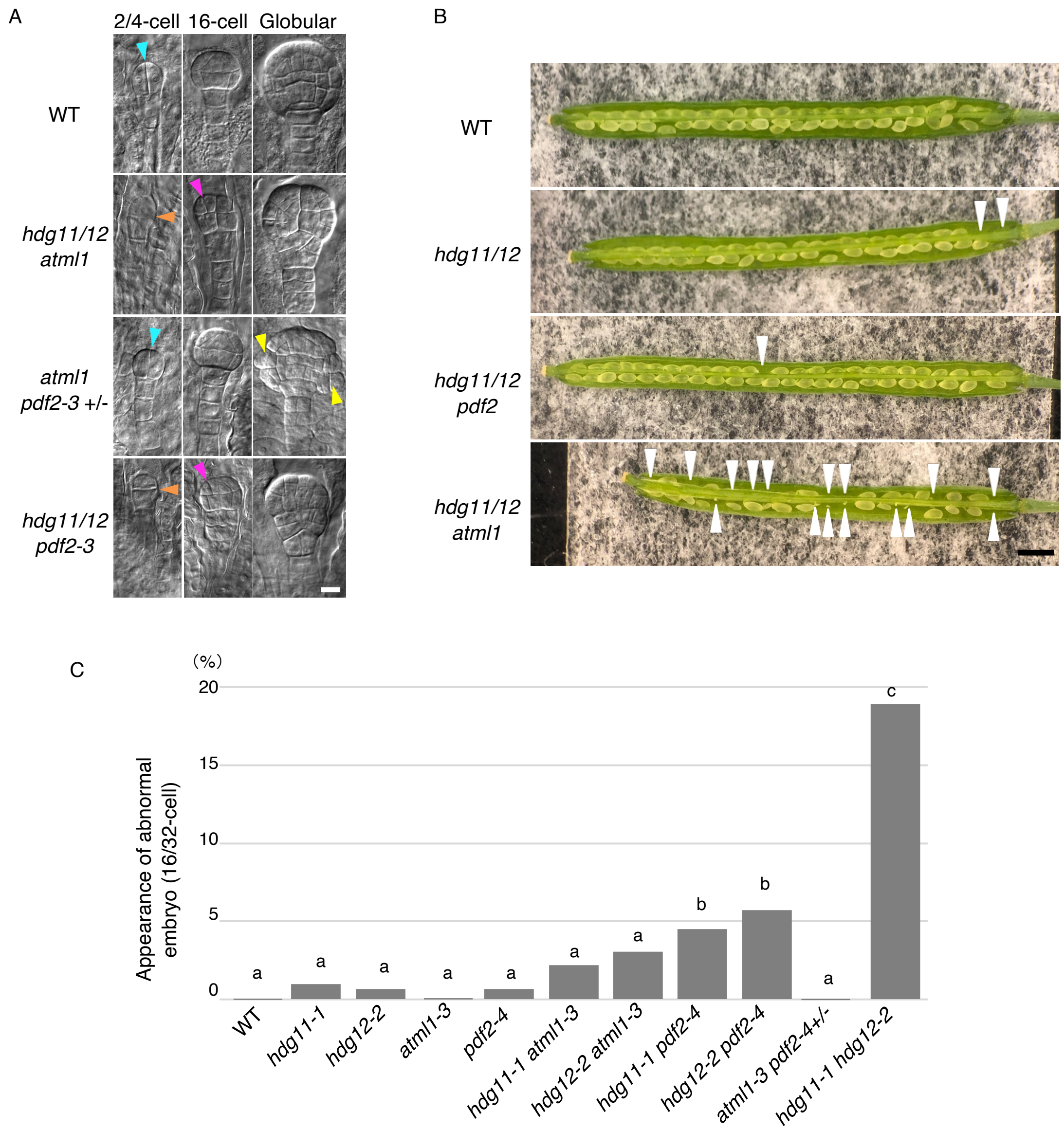


**Fig. S1.** Redundancy of *HDG11/12*, *AtML1* and *PDF2*.

(A) DIC images of cleared embryos. Genotypes and embryo stages are indicated. Cyan and orange arrowheads at 2/4-cell stage point the proper (vertical) and abnormal (transverse) cell division planes of the apical cells, respectively. Magenta and yellow arrowheads show the abnormal (anticlinal) cell division plane in proembryo and the swelled epidermal cells, respectively. (B) Opened siliques of indicated genotypes. White arrowheads point the unfertilized ovules, which left void spots in the silique. (C) Appearance of abnormal embryos, whose cell division patterns are different from wild type (WT) at 16/32-cell stages in the various combination mutants. The letters on the graph indicate significant differences determined by the Tukey–Kramer test; $P < 0.05$; n ≥ 101.

Scale bars: 10 μm (A) and 1 mm (B).

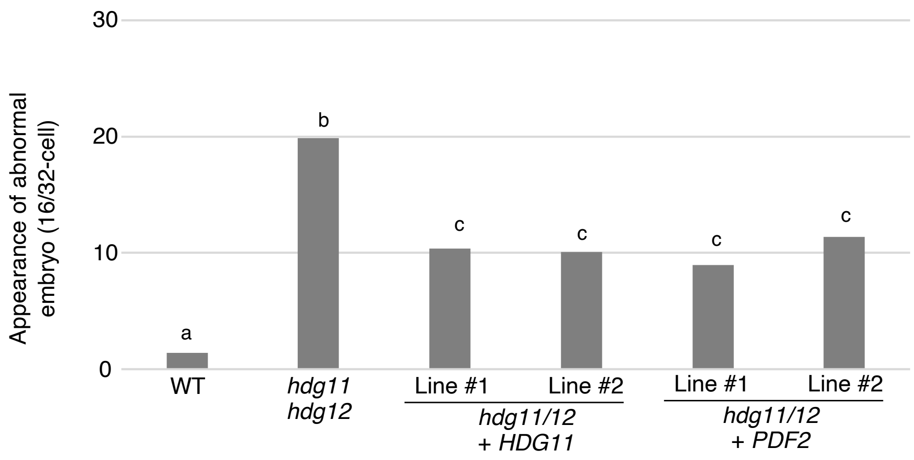


**Fig. S2.** Protein swap of HDG11 and PDF2.

Appearance of abnormal embryos, whose cell division patterns are different from WT at 16/32-cell stages in the *hdg11/12* mutants carrying no, HDG11 (*HDG11p::HDG11-YFP*), or PDF2 (*HDG11p::PDF2-YFP*) transgenes. The data of two independent lines for each transgene are shown (line #1 and 2). The letters on the graph indicate significant differences determined by the Tukey–Kramer test; $P < 0.05$; $n \geq 106$.

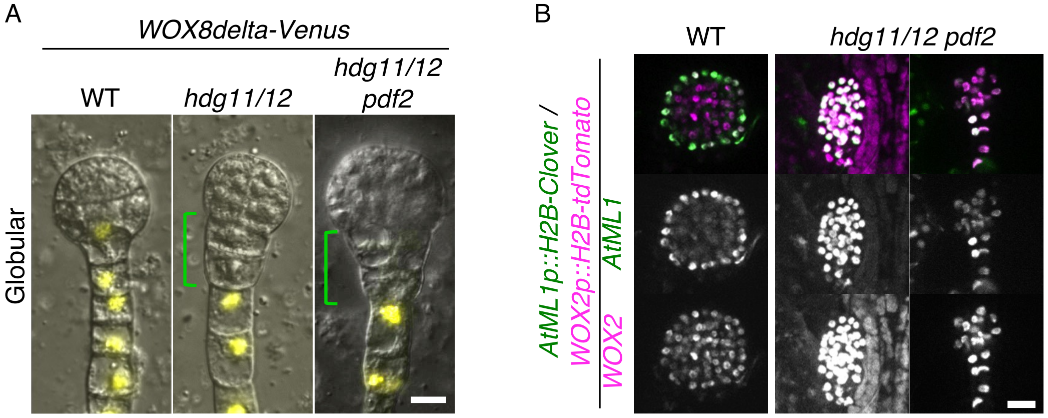


**Fig. S3.** Roles of *HDG11/12* and *PDF2* on apical-basal and radial patterns.

(A) Epi-fluorescent images of a suspensor-specific marker *WOX8delta-Venus* at globular stage. DIC images are merged to show embryo structures. Green rectangles indicate the top part of suspensors, which show abnormal cell division and no YFP signals. (B) Confocal images of an epidermis-specific marker *AtML1p::H2B-Clover* merged with an embryonic nuclei marker *WOX2p::H2B-tdTomato* at globular stage. Left and right panels of *hdg11/12 pdf2* show two different types of abnormal embryos. Lower panels show the enhanced images of individual fluorescence.
Scale bars: 10 μm.

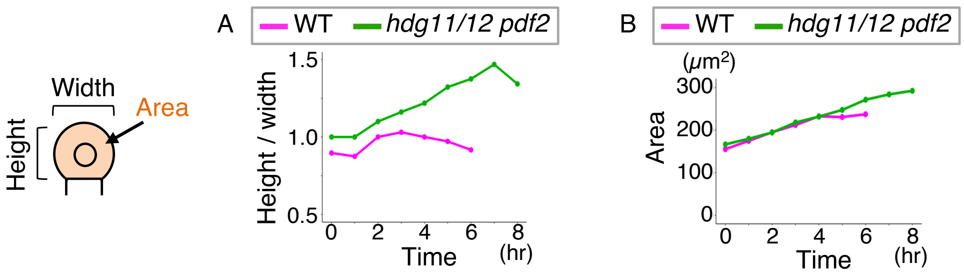


**Fig. S4.** Time-course analysis of apical cell dynamics.

(A and B) Time course of the ratio of apical cell height divided by width (height/width; A) and apical cell area (B) of the samples shown in Fig 3A and B, as shown in the left illustration. All values were measured from 0:00 when the zygote divided until before the apical cell divided in WT (6:00) and *hdg11/12 pdf2* (8:00) at 1-hour interval.

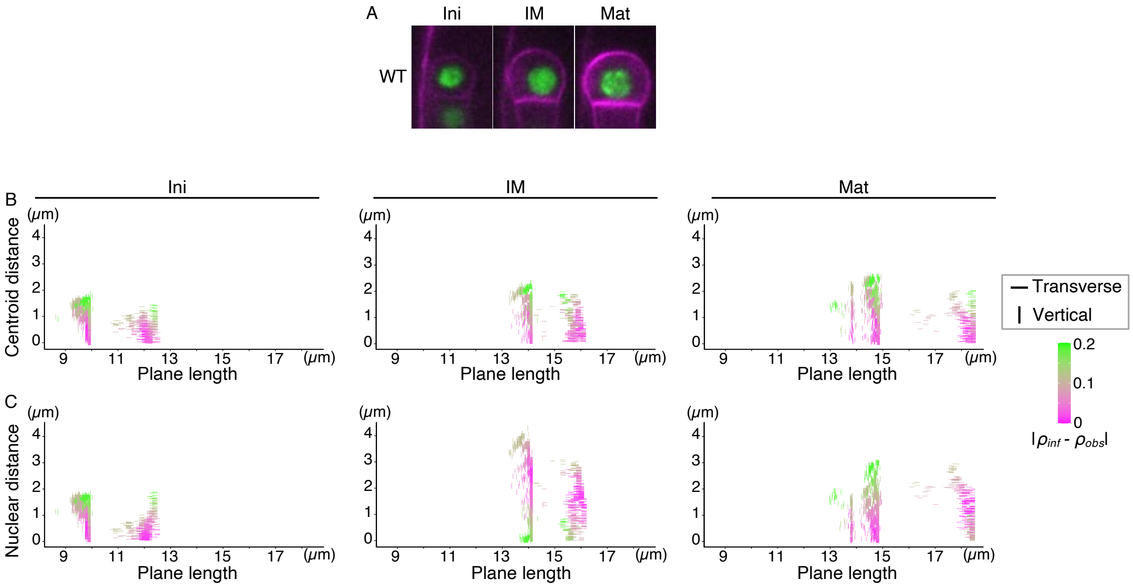


**Fig. S5.** Cellular Potts Model simulation against the apical cells at different stages in WT.
(A) Time-lapse images of WT apical cell at indicated stages, which were used for the simulation. (B and C) Simulated cell division planes. Distribution on the plane length and distance to cell centroid (B) and nuclear center (C) are shown. Simulated planes with vertical (90° ± 15) and transverse (0° ± 15) angle are plotted as vertical and transverse bars, respectively. The right color bar shows the absolute value of the difference of $\rho_{inf}$ from $\rho_{obs}$, which is set as 0. Note that the plots of Mat stage correspond to those in Fig. 4.

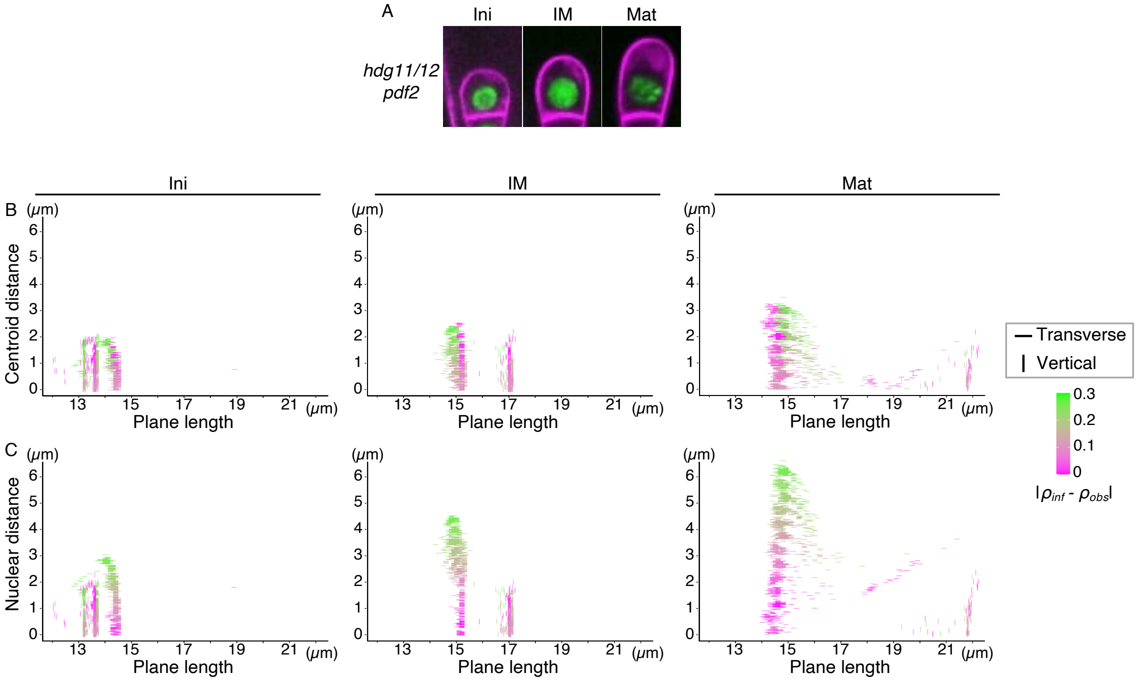


**Fig. S6.** Cellular Potts Model simulation against the apical cells at different stages in *hdg11/12 pdf2*.

(A) Time-lapse images of *hdg11/12 pdf2* apical cell at indicated stages, which were used for the simulation. (B and C) Simulated cell division planes. Distribution on the plane length and distance to cell centroid (B) and nuclear center (C) are shown. Simulated planes with vertical (90$^{\circ}$ ± 15) and transverse (0$^{\circ}$ ± 15) angle are plotted as vertical and transverse bars, respectively. The right color bar shows the absolute value of the difference of $\rho_{inf}$ from $\rho_{obs}$, which is set as 0. Note that the plots of Mat stage correspond to those in Fig. 5.

**Movie S1 (separate file).** Living dynamics from the zygote to 16-cell stage embryo in wild type (WT).

Time-lapse observation in *in vitro* cultivated WT ovules, which expressing a nuclear/PM marker, *WOX2p::H2B-GFP* and *WOX2p::LTI-tdTomato*. Numbers indicate the time (h:min), and the time when the zygote divided was set to zero. Images were obtained at 10-min intervals.
Scale bar: 10 μm.

**Movie S2 (separate file).** Living dynamics from the zygote to 16-cell stage embryo in *hdg11/12 pdf2*.

Time-lapse observation in *in vitro* cultivated *hdg11/12 pdf2* ovules, which expressing a nuclear/PM marker (*WOX2p::H2B-GFP* and *WOX2p::LTI-tdTomato*). Numbers indicate the time (h:min), and the time when the zygote divided was set to zero. Images were obtained at 10-min intervals.
Scale bar: 10 μm.

**Movie S3 (separate file).** Living dynamics from the zygote to 2-cell stage embryo in WT.

Time-lapse observation in *in vitro* cultivated WT ovules, which expressing a nuclear/PM marker, *WOX2p::H2B-GFP* and *WOX2p::LTI-tdTomato*. Numbers indicate the time (h:min), and the time when the zygote divided was set to zero. Images were obtained at 10-min intervals.
Scale bar: 10 μm.

**Movie S4 (separate file).** Living dynamics from the zygote to 2-cell stage embryo in *hdg11/12 pdf2*.

Time-lapse observation in *in vitro* cultivated *hdg11/12 pdf2* ovules, which expressing a nuclear/PM marker (*WOX2p::H2B-GFP* and *WOX2p::LTI-tdTomato*). Numbers indicate the time (h:min), and the time when the zygote divided was set to zero. Images were obtained at 10-min intervals.
Scale bar: 10 μm.
